# To touch or to be touched? Comparing pleasantness of vicarious execution and reception of interpersonal touch

**DOI:** 10.1101/2023.10.08.561444

**Authors:** Niccolò Butti, Cosimo Urgesi, Francis P. McGlone, Viola Oldrati, Rosario Montirosso, Valentina Cazzato

**Author notes:** Correspondence to: Dr Valentina Cazzato, School of Psychology, Faculty of Health, Liverpool John Moores University, UK.

## Abstract

Unmyelinated C-Tactile (CT) fibres are activated by caress-like touch, eliciting a pleasant feeling that decreases for static and faster stroking. Previous studies documented this effect also for vicarious touch, hypothesising ‘mirror-neurons’ mechanisms driving the perception and appreciation of observed interpersonal touch. Notably, less is known about appreciation of vicarious execution of touch, that is as referred to the one giving gentle touch. To address this issue, 53 healthy participants were asked to view and rate a series of videoclips displaying an individual being touched by another on hairy (i.e., hand dorsum) or glabrous (i.e., palm) skin sites, with touch being delivered at CT-optimal (5 cm/s) or non-CT optimal velocities (0 cm/s or 30 cm/s). Following the observation of each clip, participants were asked to complete self- and other-directed ratings of the pleasantness of touch for both executer (toucher-referred) and receiver (touchee-referred). Consistent with the CT fibres properties, for both self- and other-directed judgements of touch execution and reception, participants provided higher ratings for touch delivered at CT-optimal than other velocities, and when CT-optimal touch was delivered to the hand-dorsum compared to the palm. However, higher ratings were attributed to reception compared to execution of CT-optimal touch. Notably, individual differences in interoceptive trusting and attitude to interpersonal touch were positively correlated with, respectively, toucher- and touchee-related overall pleasantness of touch. These findings suggest that the appreciation of both toucher- and touchee-referred vicarious touch is specifically attuned to CT-optimal touch, even though they might rely on different neurocognitive mechanisms to understand affective information conveyed by interpersonal tactile interactions.

## Introduction

Interpersonal tactile interactions are pervasive in the early stages of life and play a pivotal role in forming social bonds and communicating emotional information throughout the life span (Cascio et al., 2019; Montirosso & McGlone, 2020). At a somatosensory level, touch pleasantness correlates with the activity of slow-unmyelinated afferents, called C-Tactile (CT) fibres (Olausson et al., 2010; Vallbo et al., 1999). CT fibres, predominantly innervating hairy skin, respond optimally to touch delivered at velocities of 1-10 cm/s, at a temperature similar to human skin (Ackerley, 2022; Ackerley, Backlund Wasling, et al., 2014), eliciting a pleasant sensation that decreases for faster and slower touch following a typical inverted-U shaped pattern (Löken et al., 2009; McGlone et al., 2014). Through their projections to the dorsal posterior insula (Kirsch et al., 2020; Morrison, 2016), CT afferents would support the processing of affective information entailed by social touch. Accordingly, recent research has documented the functional role of the CT afferents in affect regulation and social development (Fotopoulou et al., 2022), highlighting, for instance, the role of affective touch reception in modulating parasympathetic activation (Pawling et al., 2017; Van Puyvelde et al., 2019), in reducing pain (Gursul et al., 2018; Habig et al., 2017; Liljencrantz et al., 2017), and in buffering feelings of social exclusion (Von Mohr et al., 2017).

Beyond the somatosensory effects of touch, there is evidence of affective responses also when interpersonal gentle touch is just ‘observed’. Within the framework of the ‘Embodied simulation theory’ of touch (Gallese & Ebisch, 2013), individuals can map others’ tactile events by re-using their own motor, somatosensory and visceromotor representations. As a result, this tactile mapping would allow an observer to perceive the touch as if that person was receiving the same kind of tactile stimulation, thus facilitating the understanding of how the other person is ‘feeling’ that touch (Keysers et al., 2010). In line with this, functional neuroimaging evidence reports the activation of the dorsal posterior insula during vicarious observation of CT-optimal touch (Morrison et al., 2011), suggesting a similar hedonic response to directly felt and vicarious touch experiences. Furthermore, observation of vicarious interpersonal touch is rated as more pleasant when delivered at CT-optimal compared to non-CT optimal velocities (i.e., slower or faster touch), eliciting the typical inverted-U function between vicarious pleasantness ratings and velocities (Bellard et al., 2022; Devine et al., 2020; Haggarty et al., 2021; Walker et al., 2017). These findings suggest that the human brain may be attuned to “see” CT-specific features when watching others performing interpersonal touch actions (Morrison et al., 2011).

Little is known, however, about the vicarious representation of *giving* gentle touch. Indeed, observing others’ actions triggers an inner simulation of the movements in the observer’s motor system (Fadiga et al., 1995; Gazzola & Keysers, 2009). This simulative representation includes not only movement kinematics, but also the affective valence of movement (Craighero & Mele, 2018; Finisguerra et al., 2021; Urgesi et al., 2020; Vicario et al., 2019). At a higher-order socio-affective level, affective touch may have important social meaning for both those who give and those who receive it (Schirmer, Cham, et al., 2022). Evidence suggests that affective touch has a beneficial effect also for the person delivering the stroking gesture (i.e., toucher), given that those who give touch may convey feelings of closeness and care toward a touchee, who in turn may feel bonded and safe (Debrot et al., 2021; Jakubiak & Feeney, 2017). In primates, social interactions involving social touch are of critical importance for group life, so that delivering social touch is associated with desirable individual and social benefits (Jablonski, 2021). Furthermore, stroking other’s skin is perceived as pleasant by the toucher (Triscoli et al., 2017), is associated with more positive sensory experiences when delivered at CT-optimal velocities (Gentsch et al., 2015), and is spontaneously targeted to activate CT afferents (Croy et al., 2016).

Despite these similarities between touch receiving and giving, there are also profound perceptual differences which arise from the body parts that an individual normally uses to give and receive touch. Typically, most of the touching actions, like hand-holding, cradling and embracing, are performed with the toucher’s palm contacting a touchee’s arm, shoulder, or back (Schirmer et al., 2021; Triscoli et al., 2017). Notably, while CT afferents are widely represented in the hairy skin, the palm is densely innervated by fast-conducting Aβ fibres rather than CT afferents (Watkins et al., 2021). Thus, the tactile systems might receive more readily benefits from CT signalling when activated by touch receiving than by touch execution (Schirmer, Cham, et al., 2022), suggesting a preference for reception over the hairy skin, compared to execution, with the palm, of affective touch (Triscoli et al., 2017). Additionally, giving compared to receiving touch more likely involves motor mechanisms and generates predictions about the somatosensory consequences of the stroking movements (Blakemore et al., 1998; Boehme et al., 2019). These sensorial predictions are associated with inhibitory processes that dampen the awareness of emerging somatosensory impressions (Boehme & Olausson, 2022).

Taken together, all these studies corroborate the importance of interpersonal gentle touch for positive tactile interactions and suggest that the experiences and consequences of affective touch may differ between those who give and those who receive touch. Yet, the commonalities and differences between the appreciation of touch giving and of touch receiving are unclear.

The present study sought to fill this theoretical gap by investigating toucher- and touchee-related pleasantness ratings of interpersonal CT-optimal touch. Specifically, the focus in this report is on how ‘observed’ interpersonal touch is perceived as referred to the person receiving (i.e., touchee) or giving (i.e., toucher) the touch.

In line with previous studies (Bellard et al., 2022; Walker et al., 2017), a perspective manipulation was adopted to assess differences in bottom-up and top-down processing of vicarious social touch by asking participants to provide pleasantness ratings of self-directed and other-directed touch, namely judging touch pleasantness in a first-person (i.e. pleasantness for the participant) compared to a third-person (i.e. pleasantness for the model) perspective. Indeed, the appraisal of other-directed touch might rely more on the learned expectations about the rewarding pleasantness elicited by interpersonal stroking (Peled-Avron & Woolley, 2022). That is, even though one may not like to be stroked, they should still be able to acknowledge the hedonic, universally recognised, positive value of interpersonal touch. Consistent with this view, previous studies documented higher vicarious preferences for other-directed compared to self-directed touch (Bellard et al., 2022; Walker et al., 2017). To the best of our knowledge, no studies so far has investigated the effect of perspective on vicarious social touch execution and reception. For both perspectives, participants were asked to rate the pleasantness for the partner who receives or gives gentle touch.

A second aim of the study was to explore individual differences in interpersonal touch due to childhood experiences and attitudes, and interoceptive awareness, as these might facilitate (or hinder) the experience of vicarious social touch in a third-party observer, depending on perspective-taking (Bellard et al., 2022). The negative effects of childhood neglect/abuse and later life experiences on perception of affective touch are well known (Field, 2010; Keizer et al., 2022), with a recent study reporting blunted responses to vicarious affective touch in young adults who have experienced early life adversity and consequently spent time in foster care (Devine et al., 2020). Furthermore, awareness of internal bodily states, namely the sense of interoception, may play a role in vicarious social touch, with individuals with higher levels of interoceptive awareness found to show higher responses in somatosensory areas for vicarious touch perception (Adler & Gillmeister, 2019).

In line with previous evidence, the typical inverted-U shaped pattern was expected for both toucher- and touchee-referred pleasantness ratings, with overall higher pleasantness for gentle touch delivered to the hairy skin (i.e., hand), compared to the glabrous skin (i.e., palm), and at CT-optimal, compared to non-CT optimal velocities. Importantly, given that touchers and touchees may differ in the comfort they derive from interpersonal touch, higher appreciation of affective touch reception compared to execution should be found. Finally, individual differences in social touch experiences and attitudes, as well as in interoceptive awareness, would be linearly associated with toucher- and touchee-related pleasantness ratings.

## Materials and methods

### Participants

Based on a Repeated Measure ANOVA model with two agents, two perspectives, two body sites, and three stroking velocities, an a-priori power analysis using the G*Power 3.0.10 software (Faul et al., 2007) indicated that a sample > 50 allowed detecting a large effect size (*n^2^* = 0.14), with 95 % power and alpha set at 0.05 (two tailed). Hence, 53 participants (31 females, 22 males; age mean = 28.6 years, SD = 4.8) were recruited in the study. Of these, 45 individuals participated remotely, while 8 participants completed the experiment in our laboratory at Liverpool John Moores University (LJMU). It is worth noting that, for the vicarious touch task, online collected data were reported to be comparable to those of lab-based studies (Haggarty et al., 2023), and data obtained from both modalities were analysed together in a recent work (Ali et al., 2023). Participants were recruited through posters, social media advertisements, and emails to research panel lists. Inclusion criteria were: i) having normal or corrected to normal vision (with glasses/contact lenses) ii) being right-handed, iii) having no history of or any form of neurological and psychiatric disorders, iv) having no history of or any clinical condition of chronic pain and skin diseases. Participants for the lab-based study were compensated for their time with a £5 gift voucher and granted course credits if undergraduate psychology students. All procedures were approved by the LJMU Research Ethics Committee (reference number: 22/PSY/078). Informed consent was obtained by asking all participants to tick the relative box after reading the participant information sheet. The study recruitment started on the 12^th^ of February and was completed by the 16^th^ of April 2023.

### General procedure

All participants were asked to complete a vicarious affective touch task through the E-Prime 3® software (Psychology Software Tools, Pittsburgh, PA, USA), which allowed controlling stimuli administration and randomisation. The E-Prime Go® package was used for remote data collection. Then, self-report questionnaires were administered via Qualtrics® (Provo, UT, USA). Background information (e.g., age, sex, gender) was also collected, and participants were asked to confirm that they were right-handed by completing the Edinburgh handedness inventory (Oldfield, 1971). Only right-handed individuals were recruited as the displayed touchers’ movements in the vicarious social touch task were all executed with the right hand, thus potentially triggering sensorimotor representations for the same hand (Marzoli et al., 2013). Online participants were asked to sit in a quiet room and to complete the task and questionnaires in a single session. All procedures required about 30 minutes to be completed. Upon completion of the procedure participants were debriefed about the aims and instruments of the study.

### Self- and other-directed affective touch video clips

An adapted version of a previously published task was administered (Bellard et al., 2022; Walker et al., 2017). The task consisted of 6-second-long videos of both males and females applying touch with their right hand to female and male actors. Touch was delivered with CT-optimal (5 cm/s) and non-CT optimal velocities (static: 0 cm/s, fast: 30 cm/s) on the hand-dorsum and on the palm. These two body regions were selected as they were matched in terms of size and, thus, observed movements, whereas they represented areas with different density of CT-fibres, respectively, a hairy and a glabrous skin site. Moreover, the hand is considered a body part that strangers are allowed to touch (Suvilehto et al., 2015), thus pleasantness ratings of this site should be less influenced by top-down modulations related to the toucher’s identity. After viewing each video, participants were asked a series of questions concerning touch delivering and reception, which were designed to probe expectations of how touch is perceived by oneself and by others. The questions were as follows: for the toucher, “How much would you like to touch like that?” (self-directed toucher perspective) and “How pleasant do you think that action was for the person touching?” (other-directed toucher perspective); for the touchee, “How much would you like to be touched like that?” (self-directed touchee perspective) and “How pleasant do you think that action was for the person being touched?” (other-directed touchee perspective). Participants answered through a Visual Analogue Scale (VAS) scale ranging from 0 = “Not at all” to 100 = “Extremely”, for self-directed touch, and from 0 = “Very unpleasant” to 100 = “Extremely pleasant”, for other-directed touch. Separate blocks were administered for the toucher and the touchee, and within each block, self- and other-directed touch perspectives were presented in separate blocks, for a total of 4 blocks. This way, participants only answered one type of question per block (see Table 1).

**Table 1.** Task questions for each block.

|  |  | Agent |  |
| --- | --- | --- | --- |
|  |  | Toucher | Touchee |
| Perspective | Self | *“How much would you like to touch like that?”* | *“How much would you like to be touched like that?”* |
|  | Other | *“How pleasant do you think that action was for the person touching?”* | *“How pleasant do you think that action was for the person being touched?”* |

The order of administration of the four blocks was counterbalanced amongst participants. Considering a 2 agent × 2 perspective × 2 body site × 3 velocity design, 24 videos were presented once within each block in a completely randomised fashion. These videos represented all possible combinations of biological sex of actors with body sites and velocities. Overall, across all conditions and blocks, a total of 96 (i.e., 24 videos x 4 blocks) videos was presented. Examples of video stimuli are reported in Fig. 1.

**Fig. 1.**
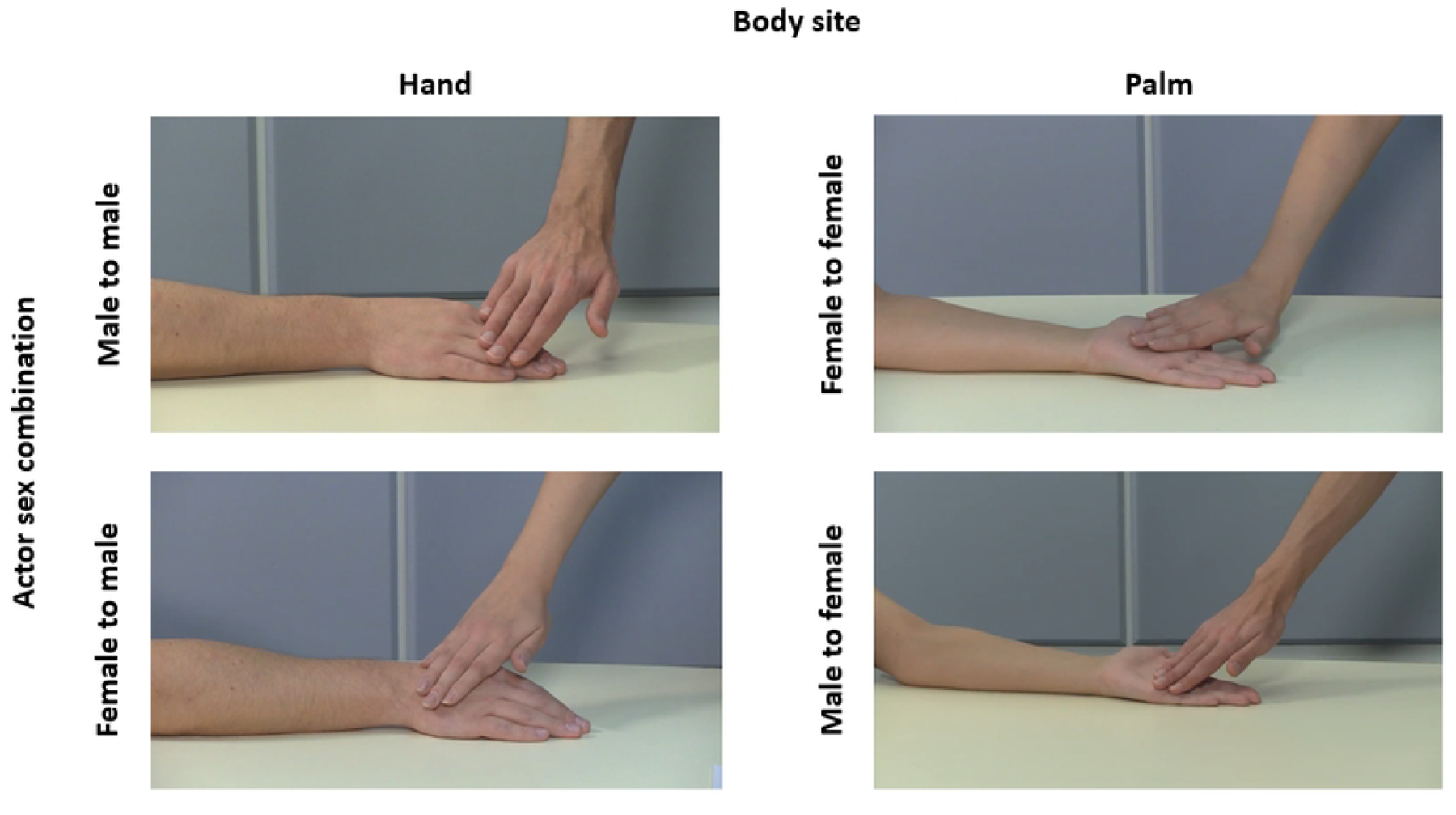
Examples of video stimuli representing touch delivered on two body sites with all combinations of biological sex of actors.

### Self-report questionnaires

#### Multidimensional Assessment of Interoceptive Awareness

The Multidimensional Assessment of Interoceptive Awareness (MAIA;) is a 32-item questionnaire which investigates eight dimensions of interoceptive bodily awareness: noticing (4 items), not distracting (3 items), not worrying (3 items), attention regulation (7 items), emotional awareness (5 items), self-regulation (4 items), body Listening (3 items) and trusting (3 items). Questionnaires are answered using a 6-point Likert scale ranging from 0 = “Never” to 5 = “Always”. Each individual dimension is scored by the average of scores from questions corresponding to that subscale, with some questions being reversed scored. Good internal consistency was reported for the MAIA questionnaire (Cronbach α = 0.90) (Valenzuela-Moguillansky & Reyes-Reyes, 2015). For this study, the noticing and trusting scales were selected, as these variables were more congruent with the hypothesis that the more a person is aware of the own bodily signals, the more this person may feel the sensations elicited by observed interpersonal touch (Ebisch et al., 2011).

#### Touch Experiences and Attitudes Questionnaire

The short 37-item version of the Touch Experiences and Attitudes Questionnaire (TEAQ) (Trotter, Belovol, et al., 2018; Trotter, McGlone, et al., 2018) assesses current and childhood experiences of positive touch and an individual’s attitude towards interpersonal touch. Questions are answered using a 5-point Likert scale ranging from 1 = “Disagree strongly” to 5 = “Agree strongly”. A mean score is calculated for each of the five subscales: attitude to friend and family touch (7 items), attitude to intimate touch (10 items), childhood touch (8 items), attitude to self-care (7 items), and current intimate touch (5 items), with negatively worded questions reversed scored. The TEAQ short version was found to have a good internal consistency (Cronbach α = 0.93). In line with our hypothesis that individual experiences and attitudes towards interpersonal touch should bias pleasantness ratings of vicarious touch, the childhood touch and the attitude to friend and family touch scales were selected as variables of interest, excluding the others as they focused on stroking the one own body and intimate touch, aspects that were less compatible with the adopted videos.

#### Data handling and statistical analysis

The pleasantness ratings attributed to each condition of the vicarious touch task were inserted into a Repeated Measure (RM) ANOVA with 2 agent (toucher vs. touchee) × 2 perspective (self vs. other) × 2 body site (hand vs. palm) × 3 velocity (3 levels: 0 cm/s, 5 cm/s and 30 cm/s) as within subject variables.

In keeping with previous research (Bellard et al., 2022; Croy et al., 2016), two indexes of touch pleasantness were calculated, namely the Overall Touch Pleasantness (OPT) and the Pleasant Touch Awareness (PTA). Since agent (i.e., touch vs. touchee) and perspective (i.e., self-vs. other-directed) represented the two main manipulations of the task, the OPT and PTA were calculated separately for agent and perspective. Indeed, for each of the four conditions (i.e., toucher, touchee, self, other) the pleasantness ratings across the different conditions of the other variable (i.e., agent, perspective) were collapsed into a single value, regardless of the two body sites (i.e., palm, hand) while considering separately the three different velocities. As an example, for the toucher condition, pleasantness ratings in the other- and self-directed blocks for both the hand and the palm were collapsed, obtaining an averaged value of the pleasantness ratings expressed for the toucher separately for static, CT-optimal and fast touch. Then, the OPT was computed as the average rating across the three velocities, thus representing an index of individual pleasantness for interpersonal touch that is not CT-specific. As a proxy of individual preference towards CT-optimal velocity, the PTA index was calculated using the following formula: (CT-optimal – non-CT optimal fast velocity)/OPT. Spearman’s *r* correlations were run between both OPT and PTA corresponding to the four different conditions, and the selected scales of the questionnaires, namely, noticing and trusting for the MAIA, and attitude to friend and family touch and childhood touch for the TEAQ.

All analyses were performed with Statistica 8.0 (Statsoft, Tulsa, OK), and data were reported as Mean ± Standard Error of the Mean (SEM). The significance threshold was set at *p* < 0.05 for all effects. Significant interactions were analysed with Duncan’s post-hoc test correction for multiple comparisons, which allows testing effects of different size in the same design (Duncan, 1955; McHugh, 2011). For the correlations, the Bonferroni correction was adopted to adjust the standard *p*-value according to the number of comparisons (corrected *p* = 0.013). Effect sizes were estimated and reported as partial eta squared (*n^2^*) for ANOVA designs, adopting conventional cut-off of 0.01, 0.06, and 0.14 for small, medium, and large effect sizes, respectively, and as Cohen’s d for pairwise comparisons, adopting conventional cut-off of 0.2, 0.5, and 0.8 for small, medium, and large effect sizes, respectively, (Cohen, 2013).

## Results

The analysis yielded significant main effects of perspective (self: 48.15 ± 1.32, other: 53.16 ± 0.98; *F_1,52_* = 39.51, *p* < 0.001, *n^2^* = 0.43) and velocity (0 cm/s: 50.70 ± 1.63, 5 cm/s: 65.75 ± 1.74, 30 cm/s: 35.52 ± 1.86; *F_2,104_*= 81.72, *p* < 0.001, *n^2^* = 0.61), which were further qualified by a significant 2-way interaction of perspective × velocity (*F_2,104_* = 5.67, *p* = 0.005, *n^2^* = 0.10). Post-hoc tests indicated that, across perspectives, CT-optimal touch was preferred than both non-CT optimal velocities, with fast stroking being judged as less pleasant than static touch (all *p* < 0.001; all Cohen’s d > 1.01). In addition, across velocities, higher ratings were attributed to other-directed compared to self-directed touch (all *p* < 0.018; Cohen’s d: static = 0.23, CT-optimal = 0.30, fast = 0.56). Moreover, a significant agent × velocity interaction (*F_2,104_* = 5.23, *p* = 0.007, *n^2^* = 0.09) revealed lower pleasantness for toucher-than touchee-referred ratings only for CT-optimal velocity (5 cm/s: 64.05 ± 1.88 vs. 67.45 ± 1.82; *p* = 0.005; Cohen’s d = 0.25), while such a difference did not emerge for static (0 cm/s: 51.09 ± 1.74 vs. 50.32 ± 1.80; *p* = 0.513; Cohen’s d = 0.06) and fast touch (30 cm/s: 36.34 ± 1.91 vs. 34.71 ± 2.08; *p* = 0.169; Cohen’s d = 0.11). Preferences for CT-optimal stroking compared to non-CT optimal velocities, and for static compared to fast touch, were detected for either agent (all *p* < 0.001; all Cohen’s d > 0.98), indicating that both toucher- and touchee-referred ratings yielded the typical inverted-U shaped pattern. The velocity × body site interaction was also significant (*F_2,104_* = 4.24, *p* = 0.017, *n^2^* = 0.08), with a preference for touch received on or delivered to the hand compared to the palm detectable only for CT-optimal velocity (5 cm/s: 66.87 ± 1.61 vs. 64.64 ± 2.13; *p* = 0.008; Cohen’s d = 0.16), and no differences between body sites for static (0 cm/s: 50.12 ± 1.78 vs. 51.28 ± 1.65; *p* = 0.162; Cohen’s d = 0.09) and fast touch (30 cm/s: 35.78 ± 1.85 vs. 35.27 ± 2.00; *p* = 0.539; Cohen’s d = 0.04). Neither the 3-way agent × velocity × body site nor any other main and interaction effects were significant (all *F* < 1.87, all *p* < 0.159). The pleasantness ratings attributed to each condition and the significant two-way interactions are represented in Fig. 2.

**Fig. 2.**
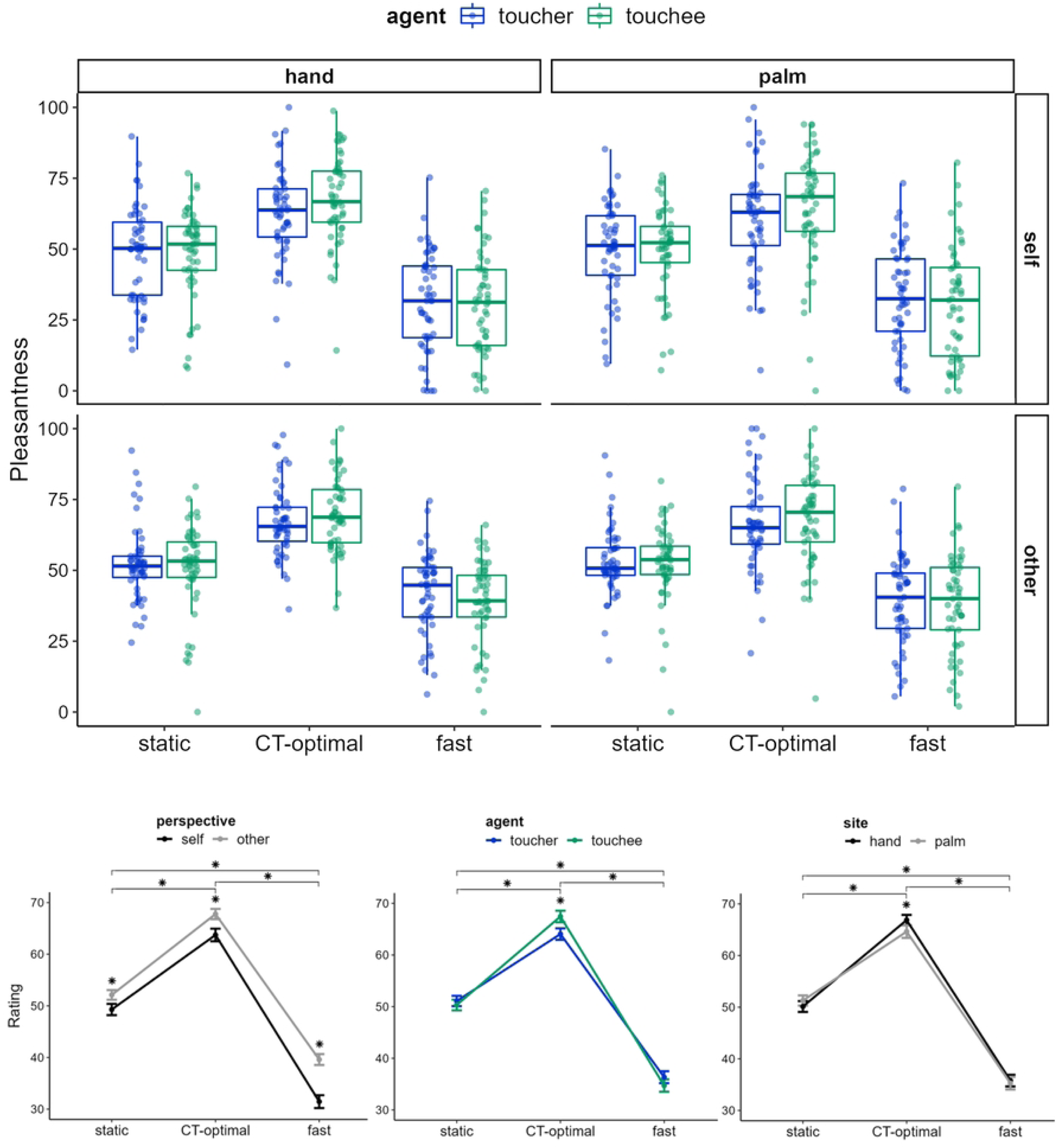
Boxplot of pleasantness ratings in the vicarious touch task and line graphs of the two-way significant interaction effects. Dots represent observations; asterisks indicate the velocity level at which the interaction effects were significant.

To sum up, for both self-and other-directed ratings of touch execution and reception, participants expressed higher pleasantness for touch delivered at CT-optimal vs. non-CT optimal velocities, and when CT-optimal touch was delivered to the hand-dorsum compared to the palm. Furthermore, higher pleasantness was attributed to touchee-compared to toucher-referred ratings only for CT-optimal touch velocities (5 cm/s). Finally, the other-directed pleasantness ratings were higher than the self-directed ones across all conditions.

### Correlation analyses

For the OTP indexes, higher scores obtained at the MAIA trusting scale were associated with higher pleasantness for other-(*r* = 0.41, *p* = 0.002) and toucher-referred (*r* = 0.36, *p* = 0.008) ratings. These indicated that the more participants trusted and felt as a safe place their own body, the higher they tended to appraise the touch pleasantness for another person, and for the person delivering the touch. In a similar vein, a more positive attitude towards friend and family touch correlated with higher ratings for other-directed (*r* = 0.36, *p* = 0.009) and touchee-referred (*r* = 0.39, *p* = 0.004) ratings. Namely, the more positive participants’ attitude to receive interpersonal touch by family and friends, the more they judged positively the observed stroking for another person, and for the person receiving the touch. All other correlations were non-significant (all *r* < 0.34, all *p* > 0.013). The significant correlations are reported in Fig. 3.

**Fig. 3.**
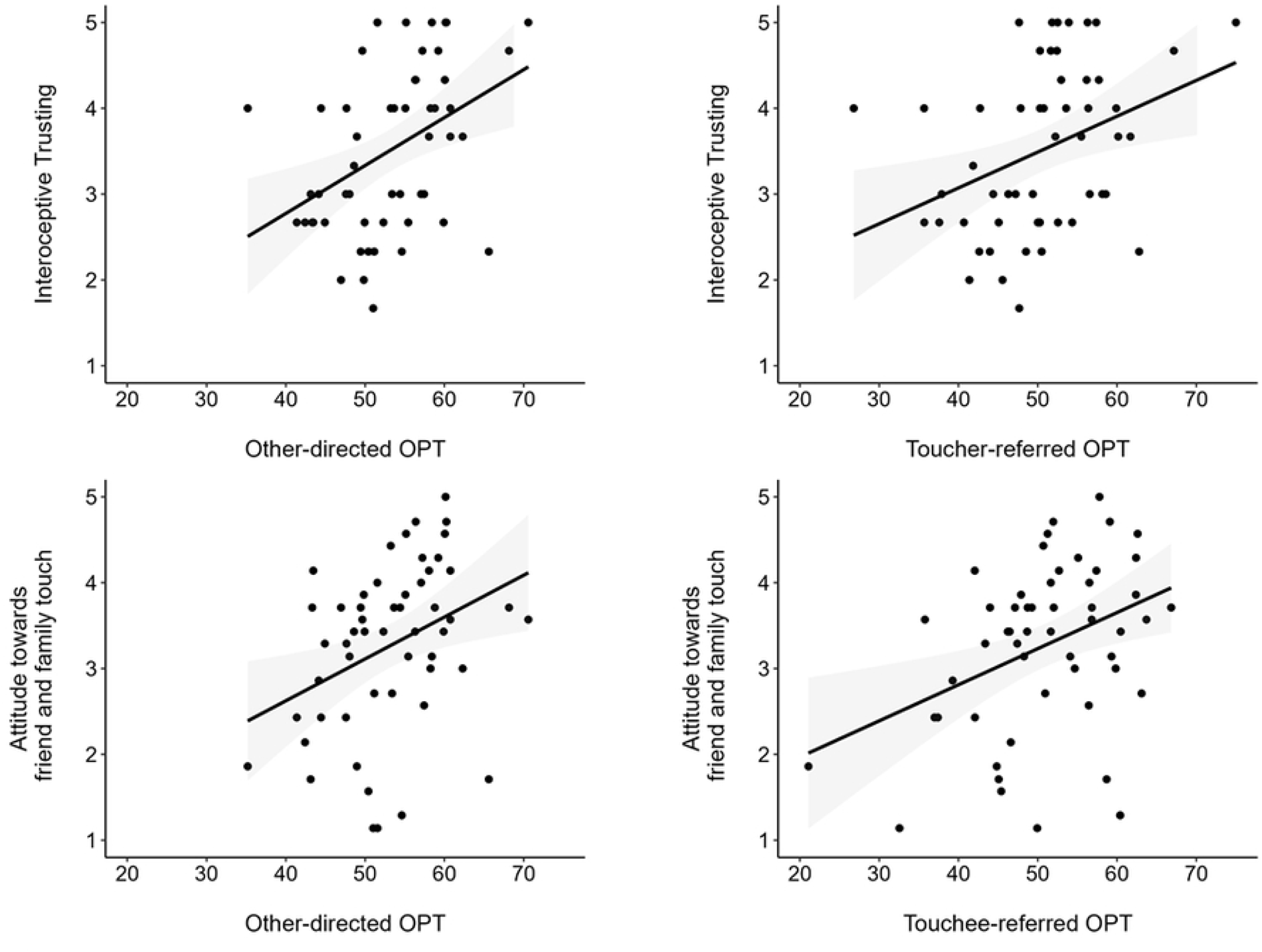
Scatter plots representing the significant correlations between OPT indexes and the questionnaire scales. Dots and represent observations; the shaded grey areas represent SE.

No significant correlations emerged between PTA and any of the questionnaire scales (all *r* < 0.24, all *p* > 0.084).

## Discussion

Previous research has found that reception of touch delivered at CT-optimal, compared to slower of faster stroking velocities, is judged as more pleasant, both when the participants directly receive the touch and when they see another person receiving it (Bellard et al., 2022; Devine et al., 2020; Haggarty et al., 2021; Walker et al., 2017). This documents an inverted-U function for the relation between pleasantness and stroking velocities for both real and vicarious touch reception. The present study aimed to compare the pleasantness ratings of vicarious touch execution and reception. Participants rated as more pleasant a touch delivered at CT-optimal than non-optimal stroking velocities when they had to embody either the partner receiving the touch or the partner delivering it. To the best of our knowledge, this is the first study documenting the typical inverted-U function characterising pleasantness ratings and velocities not only for vicarious reception (Haggarty et al., 2023), but also for vicarious delivering of interpersonal touch. These findings are in line with the hypothesis that humans are inherently ‘wired’ to receive and deliver affective touch (Croy et al., 2016; Schirmer, Cham, et al., 2022), as well as to distinguish CT-optimal stroking when observing the reception and delivery of interpersonal touch (Morrison et al., 2011), suggesting that vicarious perception of others both receiving and giving interpersonal touch is critical for adaptive behaviour in social contexts (Peled-Avron & Woolley, 2022).

The result of a CT-specificity for pleasantness ratings of vicarious delivery of touch suggests that observation of delivery of affective touch may activate embodied motor simulation of the stroking gesture, which would help individuals with understanding what kind of tactile stimulation may be more appropriate to match the touchee’s needs (Kirsch et al., 2018). Accordingly, a modulation of the Mu and Rolandic EEG rhythms, which are considered neural markers of sensorimotor simulation, was reported during vicarious touch perception (Addabbo et al., 2020; Peled-Avron et al., 2016; Schirmer & McGlone, 2019) and when participants had to carry out a consoling touch on the partner (Peled-Avron et al., 2018). However, it is less clear whether and how our motor system might map CT-specific features, such as the stroking velocity and the different CT-innervation of hairy and glabrous skin sites, during observation of interpersonal touch. For instance, it was reported that receiving or passively observing touch events does not strongly activate the motor cortex, and that the motor cortex does not represent the social nature of tactile interactions (Lee Masson et al., 2018). In this vein, our result of a CT-specificity for pleasantness ratings of vicarious delivery of touch might thus depend more on the top-down expectations that affective touch would entail positive values also for the toucher (Schirmer, Cham, et al., 2022), rather than reflecting an enhanced motor simulation for CT-optimal compared to non-CT optimal velocities. Thus, the hypothesis that our motor system may be attuned to distinguish CT features should be directly investigated in future studies, for instance testing whether CT-targeted touch modulates motor resonance processes (Naish et al., 2014).

Importantly, our findings pointed to a preference for vicarious touch reception compared to execution, which was specific for CT-optimal velocity. This preference mirrors, at a vicarious representation level, a previous study documenting that being stroked was perceived as more pleasant than stroking at CT-optimal velocities (Triscoli et al., 2017). Accordingly, such difference might depend on the scarce presence of CT fibres in the palm, so that vicarious reception would benefit more directly from CT-optimal stroking compared to vicarious execution. This view is also supported by present and previous (Walker et al., 2017) results that vicarious affective touch is perceived as more pleasant when received on or delivered to the hand-dorsum compared to the palm, according to a different distribution of CT fibres between hairy and glabrous skin (Löken et al., 2011; Watkins et al., 2021). That is, even though the inverted-U shaped pattern could be similarly elicited in glabrous and hairy skin sites (Cruciani et al., 2021), there is a functional difference between the skin typically involved in touch reception and the skin through which reaching out to touch (Ackerley, Carlsson, et al., 2014; Ackerley, Saar, et al., 2014). This critical distinction, supported also by evidence of dissociable somatosensory responses to touch on hairy and glabrous skin (Schirmer, Lai, et al., 2022), might result in the CT-specific advantage for touchee-related compared to toucher-related ratings reported here.

Furthermore, as the hedonic, positive value of a touch event would be determined by the matching between its perceived purpose and the goals of the touch receiver (Sailer & Leknes, 2022), vicarious reception of affective touch would be more important for the observer than vicarious execution in order to correctly infer affective and social information conveyed by interpersonal touch. Accordingly, an increasing number of studies documented a relationship between vicarious touch reception and affective processing, with a wide network of socio-cognitive and somatosensory areas contributing to understanding and, more importantly, feeling the observed touch (Bolognini et al., 2011, 2012; Keysers et al., 2004, 2010; Peled-Avron et al., 2019; Peled-Avron & Woolley, 2022; Rigato et al., 2019). Of note, embodied simulation in the insula and somatosensory cortices, which would subtend the “feel” of the touch, is thought to capture the perceived social intentionality of stroking gestures (Ebisch et al., 2008; Lee Masson et al., 2018; Morrison et al., 2011; Schirmer & McGlone, 2019), a mechanism that would help us to “resonate” with other’s affective experience of being touched (Lee Masson et al., 2019; Peled-Avron et al., 2016; Schaefer et al., 2012).

Our correlation results partially support the hypothesis that not only different somatosensory processes, but also diverse socio-affective mechanisms might underlie vicarious reception and execution of touch. The positive correlation between attitudes towards friend and family touch and touchee-referred ratings is in line with the rewarding value of social touch for the touchee (Morrison et al., 2010), so that the more individuals are keen to engage in interpersonal tactile interactions with people close to them, the more touch reception gains a positive salience, and thus is rated as more pleasant (Suvilehto et al., 2015). In keeping with a previous study showing that interoceptive signals may affect touch delivery (Bytomski et al., 2020), higher toucher-referred ratings were also reported by individuals with higher trust in their own-body signals. Indeed, the way a person perceives the own bodily signals plays an active role in driving behaviour and predicting the consequences of one’s actions (Seth & Friston, 2016). Hence, higher interoceptive awareness might help to understand which touch stimulation may match the touchee’s expectations, leading to increased pleasantness judgments of interpersonal touch. It should be noted, however, that these correlations were not specific for CT-optimal velocity, since only the OTP (collapsing pleasantness rating across velocities), but not the PTA (reflecting selective pleasantness for CT-optimal vs. non optimal velocities) index showed significant correlations. Thus, the associations might be related to general top-down mechanisms influencing vicarious touch perception rather than to the selective mapping of touch features related to the physiology of CT afferents (Peled-Avron & Woolley, 2022).

Importantly, as expected from previous studies (Bellard et al., 2022; Walker et al., 2017), a general advantage for other-compared to self-directed pleasantness ratings of touch emerged for both toucher- and touchee-related judgments. This is in line with the idea that humans can rely on the learned value of social touch when judging touch pleasantness for a third person (Peled-Avron & Woolley, 2022). Notably, other-directed ratings were significantly associated with both attitudes towards family and friend touch and interoceptive trusting. These results confirm that, even for vicarious delivery of touch, third-person perspective judgements are influenced by top-down expectations on the pleasantness of interpersonal tactile stimulations (Ellingsen et al., 2016). However, the difference of pleasantness ratings according to the perspective might be due to the stimuli used. Indeed, only videos viewed from a third-person perspective were presented, which might have facilitated the embodiment and relative pleasantness judgement of other-vs. self-related states.

Our study had limitations that should be carefully considered. First, although the perspective manipulation was in accordance with previous studies (Bellard et al., 2022; Walker et al., 2017) and aimed to assess differences in self-and other-directed judgements, videos were not presented from a first-person viewing perspective. Future research should consider to add videos recorded from a first-person viewing perspective, so that participants would be shown the touch event as if they were the toucher or the touch receiver, thus likely facilitating embodied simulations of the observed action (Kessler & Thomson, 2010). Furthermore, the task was designed to avoid giving any clues about the context in order to focus on CT-related manipulations (i.e., velocity, body site). However, this way it did not consider contextual and social factors that may affect vicarious touch perception (Sailer & Leknes, 2022). Finally, despite educated speculations about different neurocognitive mechanisms for vicarious touch reception and execution were advanced, this study exclusively adopted behavioural measures. Future research should investigate neurophysiological evidence of vicarious touch processing, as this topic has just started to be elucidated (Peled-Avron & Woolley, 2022).

Limitations notwithstanding, this study is the first to provide evidence of a CT-specific preference for vicarious touch execution, and it hints at different cognitive and affective mechanisms which might subserve simulation of vicarious reception and execution of social touch. These findings may pave the way for a deeper understanding of vicarious touch processing, with potential implications for psychopathological disorders showing altered touch perception and social cognition deficits. Further investigations of vicarious touch are needed to shed light on psychopathology conditions characterised by social cognition deficits and altered processing of tactile stimuli, such as autism spectrum conditions (Haggarty et al., 2020; Peled-Avron & Shamay-Tsoory, 2017), eating disorders symptomatology (Cazzato et al., 2021; Bellard et al., 2022; Crucianelli et al., 2016) and schizophrenia (Ebisch et al., 2013). A better understanding of how vicarious touch reception and execution are processed in these disorders might indeed help finding new target for rehabilitation treatments, also considering the potential use of virtual reality to provide multisensory stimulation that can shape touch processing, overcoming at the same time social anxiety often present in these conditions (Della Longa et al., 2022; Spence, 2022).

## Declarations

### Authors’ contributions

NB conceived the study. VC and CU contributed to the concept and design of the study. VO helped with implementation of the paradigm. NB performed data collection and analysed the data, under the supervision of VC and CU. The first draft of the manuscript was written by NB, with support from all authors. All authors approved the manuscript before submission.

### Data availability statement

All the relevant data are freely available from OSF at the following weblink: https://osf.io/3b9wy/?view_only=aa741b03542547b0b0b863729a47742d.

### Ethics Statement

All procedures were approved by the LJMU Research Ethics Committee (reference number: 22/PSY/078) and complied with the ethical standards of the 1964 Declaration of Helsinki.

## Acknowledgements

A special thanks goes to Adarsh Makdani for providing technical support. We would like also to thank the SomAffect research group at LJMU for supporting the project with inspiring discussion.

